# PLM-OMG: Protein Language Model-Based Ortholog Detection for Cross-Species Cell Type Mapping

**DOI:** 10.1101/2025.05.30.657127

**Authors:** Tran N. Chau, Song Li

**Affiliations:** Genetics, Bioinformatics and Computational Biology, Virginia Tech, Blacksburg, Virginia, USA; School of Plant and Environmental Sciences, Virginia Tech, Blacksburg, Virginia, USA; Department of Computer Science, Virginia Tech, Blacksburg, Virginia, USA

## Abstract

Understanding conserved and divergent cell types across plant species is essential for advancing comparative genomics and improving crop traits. Accurate and scalable ortholog detection is central to this goal, particularly in cross-species single-cell analysis. However, conventional methods are time-consuming and perform poorly with distantly related species, limiting their effectiveness. To address these limitations, we introduce PLM-OMG, a protein language model-based framework for orthogroup classification and cross-species cell type mapping. We benchmark five deep learning models including ESM2, ProGen2, ProteinBERT, ProtGPT2, and LSTM, using a curated 15-species dataset and large-scale monocot and dicot datasets from PLAZA. Transformer-based models, particularly ProtGPT2 and ESM2, achieve superior accuracy and generalization across evolutionary distances. Our results show that PLM-OMG enables scalable and reusable orthogroup detection without recomputing existing groups, significantly reducing computational overhead and highlighting its potential to transform cross-species transcriptomic analysis in plant genomics.

## 1 Introduction

Cross-species single-cell analysis is crucial for understanding both conserved and divergent cell types, and is important for transferring cell type knowledge from model species to non-model species [1]. This is particularly important for plant genomic research, as plants are the primary source of food, fuel, and fiber for human society [2]. However, most research on gene function has been conducted using the model plant species Arabidopsis thaliana, which is not directly applicable to crop production [2]. Consequently, plant biology research heavily relies on the ability to transfer knowledge from Arabidopsis to agriculturally important species such as tomato and maize [3]. A key step in this process is accurate cell type mapping, which depends on identifying shared gene families across species. The Orthologous Marker Gene groups (OMG) framework [4] supports this by identifying cell-type-specific marker genes that belong to orthogroups, which are sets of genes descended from a common ancestor and conserved across species. We have successfully demonstrated that the OMG method can map cell types across 15 diverse plant species. However, the current OMG framework relies on OrthoFinder [5], a widely used tool that infers orthogroups through all-vs-all sequence similarity searches followed by clustering and phylogenetic analysis.

While effective, OrthoFinder has practical limitations. It is restricted to a predefined set of species (e.g., 15 in the original OMG paper [4]), and adding new species requires re-running the full pipeline. This re-computation often leads to changes in orthogroup IDs, making results difficult to compare when single cell data from more species are included. Conventional tools like MMseqs2 [6] and DIAMOND [7] offer faster alternatives for pairwise protein sequence comparison. They allow new species to be matched against an existing reference set, enabling incremental updates without redefining existing orthogroups. However, these tools are often time-consuming and less accurate when comparing distantly related species due to reliance on local sequence similarity alone.

To overcome the limitations of traditional ortholog detection methods, we introduce PLM-OMG which is a protein language model-based framework for detecting Orthologous Marker Gene groups. In PLM-OMG, we benchmark state-of-the-art protein language models (PLMs) including ESM2 [8], ProGen2 [9], ProteinBERT [10], and ProtGPT2 [11] for orthogroup classification. ESM2, the largest of these models with up to 15 billion parameters, excels at predicting protein structure at atomic resolution directly from primary sequences, bypassing the need for multiple sequence alignments. Its scalability and precision make it well-suited for modeling protein relationships across diverse organisms [8]. ProGen2, trained on 280 million protein sequences from over 19,000 families, is optimized for functional protein generation using conditional tags to control properties of the output sequences [9]. ProteinBERT, inspired by the BERT architecture, combines bidirectional language modeling with Gene Ontology (GO) annotation prediction during pretraining. Trained on more than 106 million protein sequences, it is designed for efficient protein function prediction [10]. ProtGPT2, with 738 million parameters, adopts an autoregressive decoder-only architecture for de novo protein design, generating sequences that closely resemble natural proteins in composition and structure, while also exploring novel areas of the protein sequence space [11]. These models offer a robust alternative to alignment-based tools, by capturing structural and evolutionary features, enabling a scalable and reusable framework that avoids recomputing orthogroups when new species are added. In this study, we evaluate and compare the performance of these protein language models on orthogroup classification across multiple plant datasets, including a curated 15-species dataset and large-scale monocot and dicot datasets. We further assess their generalization across sequence similarity levels and demonstrate their utility in improving cross-species cell type mapping.

## 2. Methods

### 2.1 Datasets

#### 2.1.1 15 species datasets

This dataset comprises 712,894 protein sequences from 15 plant species (including Arabidopsis thaliana, Brassica rapa, Catharanthus roseus, Fragaria vesca, Glycine max, Gossypium bickii, Gossypium hirsutum, Manihot esculenta, Medicago truncatula, Nicotiana attenuata, Oryza sativa, Populus x pyramidalis, Populus x glandulosa, Solanum lycopersicum, Zea mays), grouped into 68,071 orthologous groups using OrthoFinder. Sequence lengths range from 19 to 4,656 amino acids, with a median length of 331 (Figure 1A). While most orthogroups contain relatively few sequences, a small subset includes substantially more, highlighting the uneven distribution and variability in gene family sizes across species (Figure 1B).

**Figure 1.**
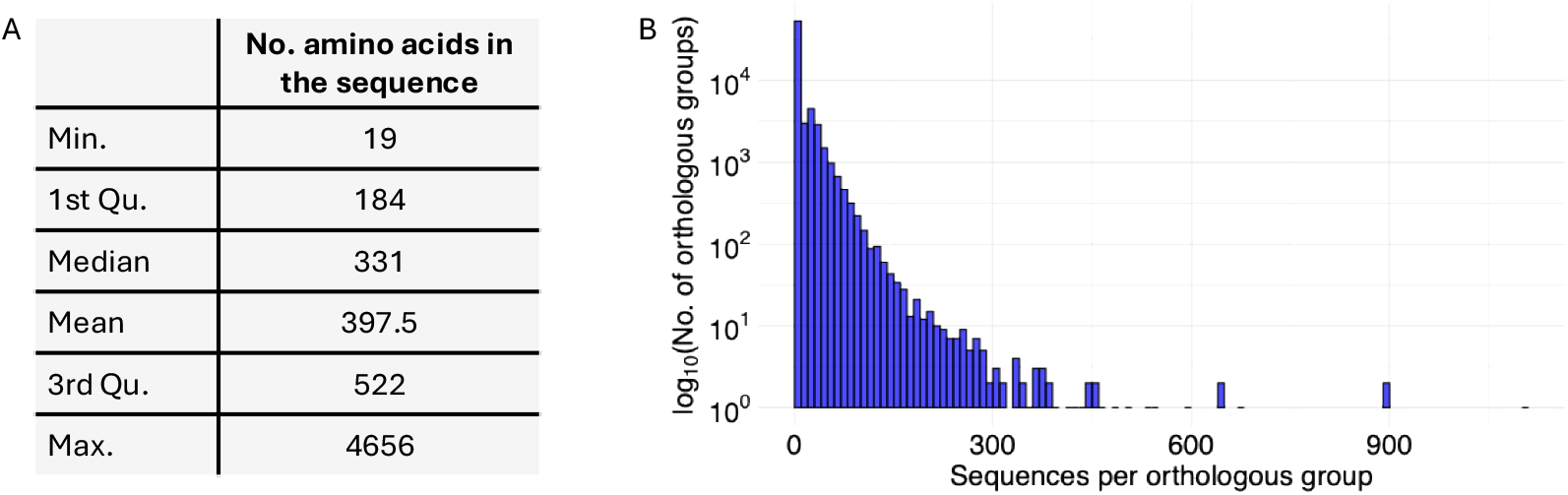
Overview of input sequences. A. Summary statistics of amino acid sequence lengths B. Distribution of the number of sequences per orthologous group (log10 scale) after quality control

#### 2.1.2 Monocots

The Monocots dataset from PLAZA 5.0 [12] integrates structural and functional annotations across 53 monocot species, a group of flowering plants characterized by having a single embryonic leaf (cotyledon). It includes over 2.25 million genes, clustered into 48,496 orthogroups.

#### 2.1.3 Dicots

The Dicots dataset, also from PLAZA 5.0 [12], integrates structural and functional annotations across 100 plant species to support large-scale comparative genomics. Dicots are flowering plants typically characterized by having two embryonic leaves (cotyledons). The dataset contains over 4.2 million genes, grouped into 68,306 orthogroups.

### 2.2 Data processing

The dataset is split by species into 80% training, 10% development, and 10% test sets. For quality control, orthologous groups with fewer than 50 genes (15-species dataset), 100 genes (monocots), and 200 genes (dicots) were excluded from the training set to ensure sufficient representation for effective protein language model training. These thresholds are chosen based on the size and complexity of each dataset to improve training efficiency and model performance, particularly for large-scale models. To maintain consistency in classification, only orthogroups present across the filtered training, development, and test sets were retained by taking their intersection. Each remaining orthogroup is then assigned a unique numerical class ID for classification tasks.

For sequence preprocessing, each amino acid is converted into a unique token to represent protein sequences in a format suitable for model input. To standardize sequence lengths, padding is applied to shorter sequences, while truncation is used for those exceeding the maximum allowed length of 512 tokens. A token-to-ID mapping is then used to assign each token a unique numerical identifier, enabling the sequences to be encoded as numeric vectors. This tokenization process is automated to streamline preprocessing and ensure consistency in data preparation for model training.

### 2.3 Models

For the ESM2 model (Table 1), protein sequences are tokenized using the pre-trained tokenizer from `facebook/esm2 t6 8M UR50D`. This encoder-only transformer-based model is configured for sequence classification, with the number of output classes matching the unique orthogroups. ESM2 can handle millions of protein sequences without reliance on multiple sequence alignments. The optimization employs AdamW (learning rate: 2e-5) and CrossEntropyLoss, and the model is trained for 10 epochs. ESM2 is particularly efficient at learning structural information due to its scale (8M to 15B parameters).

**Table 1:** Summary of Protein Language Models. Dataset column indicates the number of protein sequences used to train the models in their original reference papers.

| Model | Architecture | Parameters | Dataset | Function |
| --- | --- | --- | --- | --- |
| ESM2 | Encoder-only transformer by Meta AI | 8M to 15B | 617M | Independent of multiple sequence alignments |
| ProtGPT2 | Autoregressive decoder-only transformer | 738M | 50M | Predicts amino acid sequences, enabling novel protein generation |
| ProGen2 | Autoregressive decoder-only transformer | 151M to 6.4B | 1B | Generates proteins with properties similar to natural proteins |
| ProteinBERT | BERT architecture | 16M | 106M | Fills missing amino acids and captures both local and global patterns |
| LSTM | RNN-based model |  |  | Captures long-term dependencies for sequence analysis |

The ProGen2 model (Table 1) employs an autoregressive decoder-only transformer architecture and is trained on 280M protein sequences. Sequences are tokenized using the `hugohrban/progen2-medium`tokenizer with special tokens and right-side padding for uniform input lengths. ProGen2 generates proteins that mimic the structure and properties of natural proteins. A custom ProteinClassifier extends the base model by adding a linear classification layer, with mean pooling used to aggregate sequence embeddings for accurate orthogroup classification.

For the ProtGPT2 model (Table 1), sequences are tokenized using the `nferruz/ProtGPT2`tokenizer, which employs special tokens and padding. ProtGPT2, another autoregressive decoder-only transformer, predicts the next amino acid to enable novel protein generation. A custom ProtGPT2Classifier fine-tunes this model with a classification head that applies mean pooling to the last hidden states, producing sequence embeddings that pass through a linear layer for classification.

The ProteinBERT model (Table 1) leverages a BERT-based architecture trained on 106M protein sequences to predict missing amino acids while capturing both local and global sequence patterns.

Finally, a custom LSTM model (Table 1) for protein is developed. This model consists of an embedding layer, a bidirectional LSTM, and max-pooling for feature extraction, followed by a fully connected layer for classification. The LSTM leverages long-term dependencies in protein sequences, making it suitable for sequence analysis tasks.

## 3 Results

### 3.1 Performance and resource comparison of five deep learning models for orthogroup classification

To assess the effectiveness of various model architectures for orthogroup classification, we evaluate five deep learning methods including ProteinBERT, LSTM, ProGen2, ESM2 and ProtGPT2 using data from 15 plant species. Among these approaches, ProtGPT2 achieves the highest performance, with 0.95 accuracy, 0.94 F1 score, 0.95 precision, and 0.95 recall (Figure 2A). The ProtGPT2 training curves reveal rapid convergence with both the training and validation accuracy reaching high values within the first few epochs and maintaining stable performance throughout the 10-epoch training period (Figure 2B). ESM2 follows closely, achieving an accuracy of approximately 0.89 despite its small size of only 8 million parameters (Figure 2A, C). This lightweight model requires only 341 minutes to complete 10 epochs while maintaining strong classification performance. In contrast, ProteinBERT shows the weakest performance, with 0.36 precision, 0.30 F1 score, 0.41 precision, and 0.32 recall (Figure 2A). Despite having the second-longest training time (946 minutes), it fails to converge within 10 epochs. Training accuracy remained consistently below validation accuracy, indicating possible overfitting issues (Figure 2D). These results highlight the effectiveness of transformer-based protein language models for orthogroup classification. In particular, ProtGPT2 delivers the highest accuracy, while ESM2 offers the best trade-off between performance and computational efficiency.

**Figure 2.**
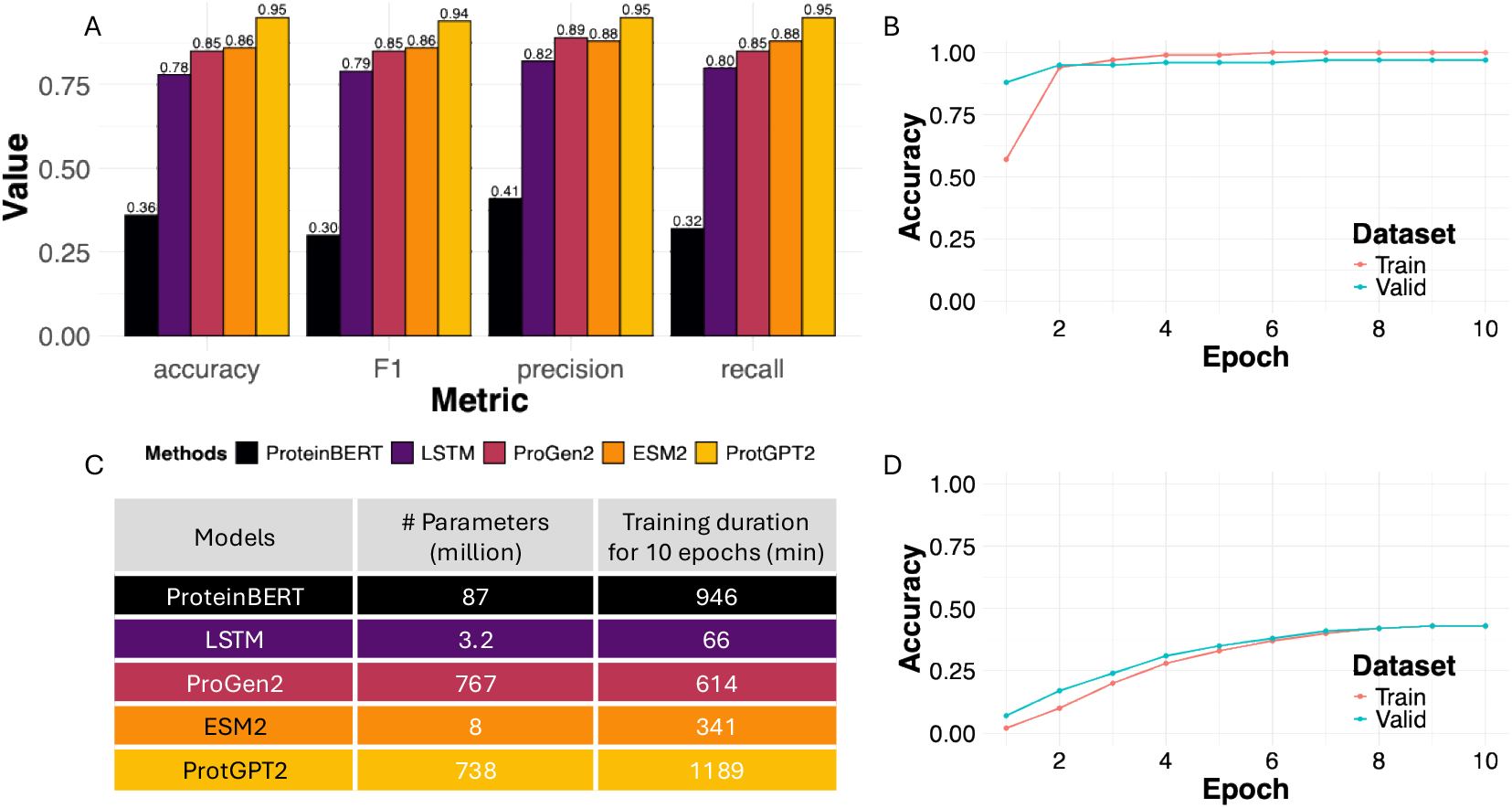
Models performance comparison. A. Model performance metrics (accuracy, F1 score, precision, and recall) across five methods B. Training and validation accuracy over epochs for ProtGPT2 C. Summary of model complexity in terms of the number of parameters and training duration for 10 epochs D. Training and validation accuracy over epochs for ProteinBERT

### 3.2 Model generalization across sequence similarity levels

To assess the generalizability of our deep learning approaches, we evaluates the performance of the model in four distinct sequence similarity groups (*<* 40%, 40-60%, 60-80%, and *>* 80% sequence identity) using MMseqs2 clustering (Figure 3). All models perform best on sequences with *>* 80% similarity, where homology signals are strongest. ProtGPT2 consistently achieves the highest accuracy and F1 score, followed by ESM2 and ProGen2, which also demonstrates strong performance (red polygons in radar plots). Even lower-performing methods such as ProteinBERT and LSTM show improved results on high similarity sequences, suggesting that all approaches benefit from clear sequence homology signal (red polygons in radar plots). As sequence similarity decreases, performance decreases across all models, with the *<* 40% similarity group posing the greatest challenge (yellow polygons). However, ProtGPT2, ESM2, and ProGen2 still maintain relatively robust performance even on these evolutionarily distant sequences. The radar plots highlight the superior generalization ability of transformer-based models, particularly ProtGPT2, across all similarity ranges (Figure 3 A-B). This indicates their strong potential for orthogroup classification in diverse and distantly related protein datasets.

**Figure 3.**
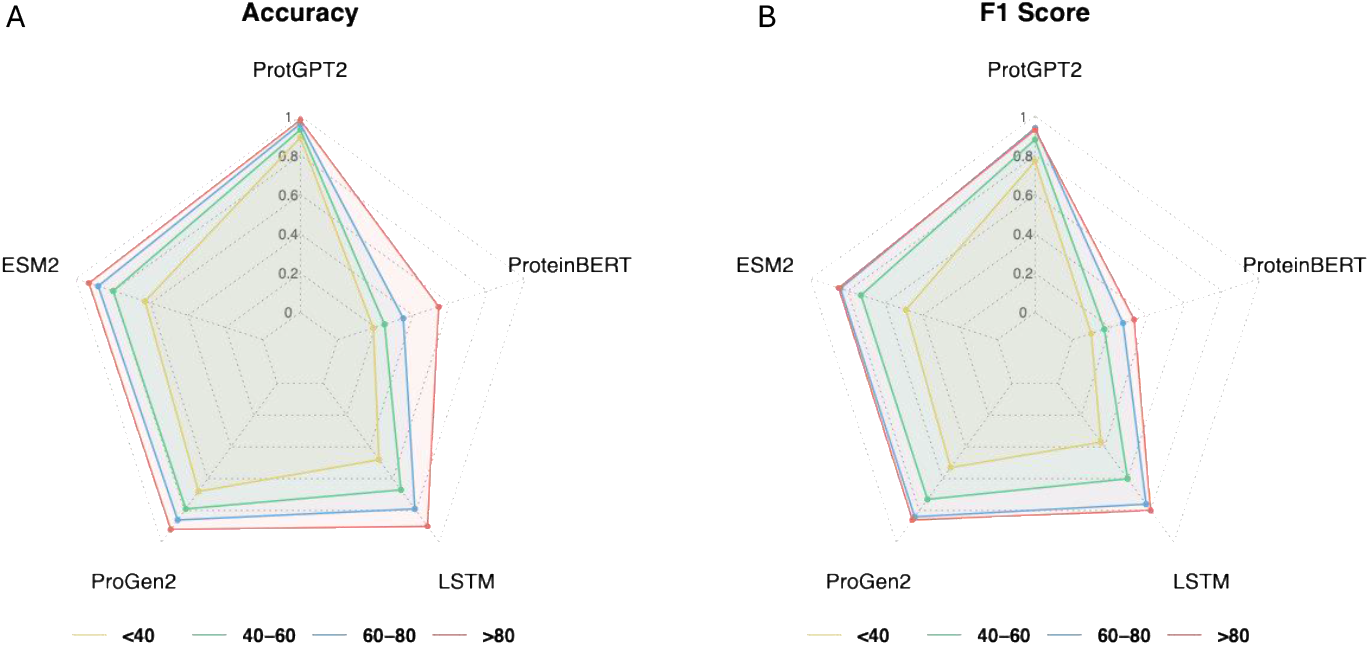
Model performance across different similarity thresholds. A-B. The accuracy and F1 score of five models (ProteinBERT, ProtGPT2, ESM2, ProGen2, and LSTM) on test orthogroups grouped by different sequence similarity thresholds

### 3.3 Robustness assessment on large-scale plant datasets

To assess the robustness and scalability of our deep learning approaches, we evaluate model performance on two comprehensive plant datasets: Monocots and Dicots Plaza 5.0. ProteinBERT performs better on the Dicots dataset than on the Monocots dataset but overall shows limited predictive ability, with consistently lower performance compared to the other models (Figure 4). ProtGPT2 demonstrates outstanding consistency, achieving the highest performance across all metrics including accuracy, precision, recall, and F1 score on both monocot and dicot datasets. This stability underscores its strong generalization capabilities across divergent plant lineages. ESM2, ProGen2, and LSTM also maintain high and consistent performance between the two datasets, further supporting their robustness (Figure 4). These findings confirm that transformer-based protein language models, particularly ProtGPT2, ESM2, and ProGen2, are highly robust for large-scale orthogroup classification.

**Figure 4.**
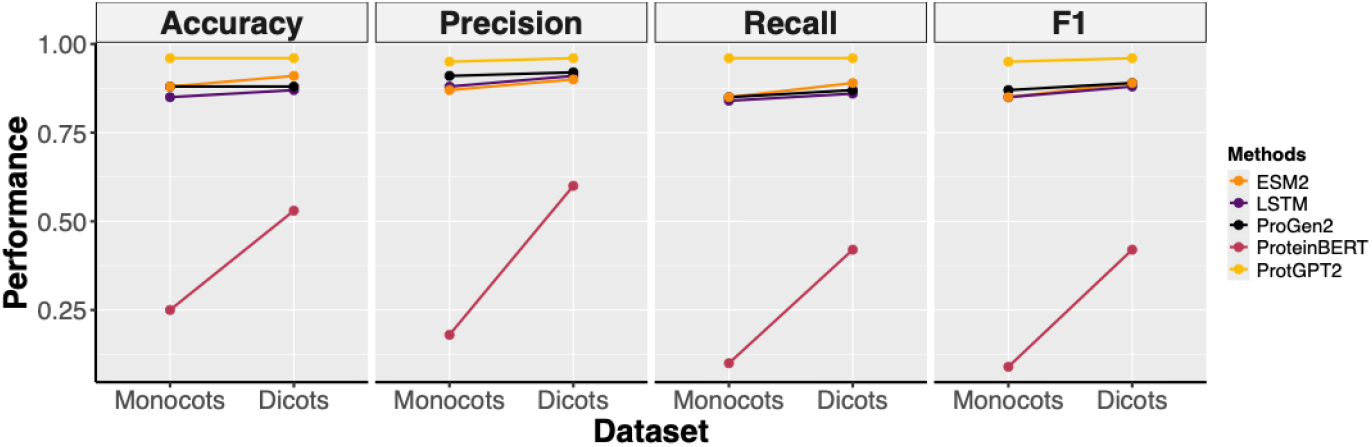
Performance comparison of protein classification models across monocot and dicot datasets. The accuracy, precision, recall, and F1 score of five models (ESM2, LSTM, ProteinBERT, ProGen2, and ProtGPT2) on monocot and dicot test sets. ProtGPT2 outperforms others across all metrics, while ProteinBERT shows poor performance on both datasets.

### 3.4 Cross-species cell type mapping

To evaluate the effectiveness of orthogroup-based methods for cross-species cell type annotation, we compare OrthoFinder and ESM2 within the OMG (Orthologous Marker Gene) framework.

ESM2 is selected because the model has fewer parameters and the performance is comparable to the ProtGPT2. The OMG framework leverages single-cell transcriptomic data by identifying marker orthogroups which groups of genes shared across species and specific to certain cell types to match cell types between species. OrthoFinder is a widely used tool that clusters genes into orthogroups based on sequence similarity and phylogenetic inference, serving as a conventional baseline for ortholog detection. Using both the traditional OrthoFinder approach and the protein language model ESM2, we map root cell types between Tomato and Arabidopsis (Figure 5A–B) and between Maize and Arabidopsis (Figure 5C–D), demonstrating the potential of PLM-OMG in transferring cell type annotations across plant species.

**Figure 5.**
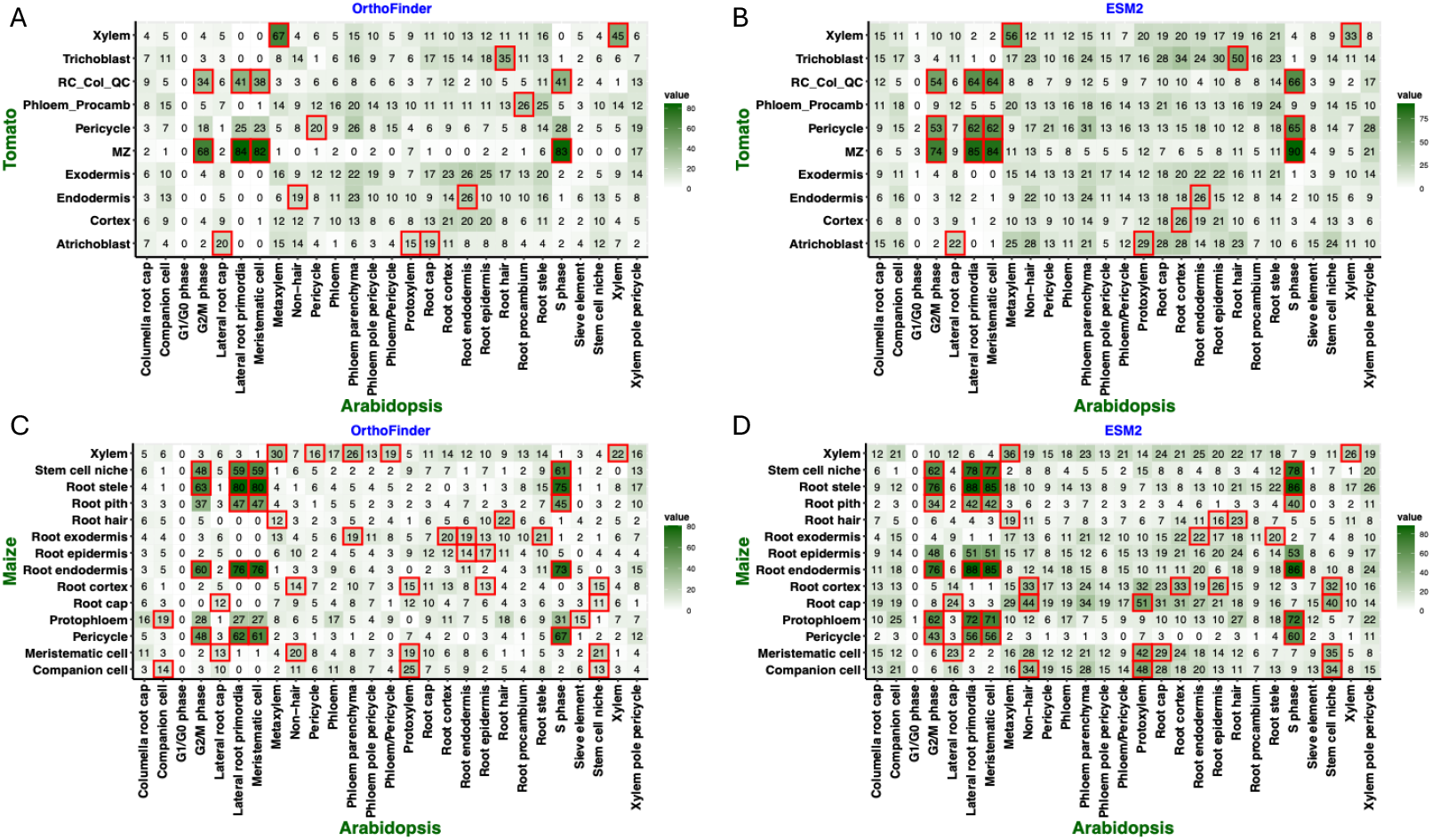
Cross-species cell type mapping using orthologous marker genes (OMGs) (A–B) Tomato vs. Arabidopsis mappings using OrthoFinder (A) and ESM2 (B). (C–D) Maize vs. Arabidopsis mappings using OrthoFinder (C) and ESM2 (D). Color intensity reflects the number of shared OMGs between cell type pairs: darker cells indicate a higher number of common marker orthogroups, while lighter cells indicate fewer. Red boxes highlight statistically significant overlaps, as determined by Fisher’s exact test (FDR *<* 0.01).

In the Tomato–Arabidopsis comparison, both methods identified biologically meaningful cell type correspondences, with 18 pairs detected by OrthoFinder and 19 pairs by ESM2. OrthoFinder matches 14 specific cell type pairs between the species, while ESM2 matches 13. However, ESM2 produces stronger and more precise signals, particularly for key cell types such as trichoblasts (root hair cells), the meristematic zone (MZ), and the cortex, with consistently higher number of conserved OMGs detected as compared to OrthoFinder based results. ESM2 also reveals distinct correspondence patterns for specialized cell types that were less evident in the OrthoFinder results. The ESM2 heatmap exhibits higher maximum intensities and sharper contrast, reflecting more accurate orthogroup assignments.

In the Maize-Arabidopsis comparison, both methods identify 11 corresponding cell type pairs, with 48 significant overlaps detected by OrthoFinder and 46 by ESM2. Despite the similar numbers, ESM2 detects more conserved orthologous marker genes for key tissues such as xylem, root hair, cortex, and root cap, suggesting improved resolution in detecting conserved gene expression profiles. These results demonstrate that ESM2 provides greater resolution and accuracy in identifying conserved cell types across species. They highlight the promise of transformer-based protein language models for improving cross-species cellular annotation and advancing comparative plant genomics.

## 4 Discussion

This PLM-OMG study demonstrates the effectiveness of protein language models (PLMs) for orthogroup classification and cross-species cell type mapping in plants. By benchmarking state-of-the-art PLMs across diverse datasets, we show that these models generalize well across evolutionary distances and outperform traditional alignment-based methods. A key advantage of this approach is its scalability and reusability, as PLMs do not require orthogroup recomputation when new species are added, making them well-suited for ongoing comparative genomics studies.

Despite these benefits, training and fine-tuning large PLMs require substantial computational resources, including high-memory GPUs. The training process is also time-consuming, which may limit accessibility for some research groups.

Data imbalance presents another challenge. To ensure sufficient representation, orthogroups with too few sequences were excluded from training. In our application of identifying orthologous marker genes, this is not a significant issue because a sufficient number of conserved genes across species is a prerequisite for defining cross-species cell type marker genes. However, ignoring orthogroups with small number of sequences may also lead to the loss of biologically meaningful information.

Moreover, to maintain consistency during training, only orthogroups present in the training, development, and test sets are used. Although this filtering step ensures fair evaluation, it reduces the diversity of orthogroups and may limit the model’s ability to handle unseen gene families. In practice, future datasets may contain orthogroups that are not represented in the training data, which could affect the accuracy of cross-species cell type comparisons. To mitigate this limitation, we extract the predicted probability for each protein’s orthogroup assignment and apply a confidence threshold. Genes with low prediction confidence are excluded, helping to reduce noise and improve the reliability of cross-species cell type mapping.

## Supporting information

Scripts

## 5 Code and Data availability

The scripts can be found at out GitHub repository (https://github.com/ct-tranchau/PLM-OMG). The processed data are deposited in Zenodo (https://zenodo.org/records/15507407).

## Notes

https://zenodo.org/records/15507407

### Competing Interest Statement

The authors have declared no competing interest.

### Funding

- National Science Foundation- Plant Genome Research Program: 2344169

