## Supplementary material for "PLM-OMG: Protein Language Model-Based Ortholog Detection for Cross-Species Cell Type Mapping": Scripts: esm2_15.py

```
import os
import pandas as pd
import torch
from torch.utils.data import Dataset, DataLoader
from transformers import AutoTokenizer, AutoModelForSequenceClassification
from torch.nn import CrossEntropyLoss
from torch.optim import AdamW
from sklearn.metrics import accuracy_score, precision_score, recall_score, f1_score, classification_report
import matplotlib.pyplot as plt

# Reproducibility
torch.manual_seed(42)

# Load datasets
train_data = pd.read_csv("/projects/intro2gds/LLMOMG/Fasta/train.csv", dtype={"seq": str})
test_data = pd.read_csv("/projects/intro2gds/LLMOMG/Fasta/test.csv", dtype={"seq": str})
test_data_lt40 = pd.read_csv("/projects/intro2gds/LLMOMG/Fasta/bin_lt40.csv", dtype={"seq": str})
test_data_40_60 = pd.read_csv("/projects/intro2gds/LLMOMG/Fasta/bin_40_60.csv", dtype={"seq": str})
test_data_60_80 = pd.read_csv("/projects/intro2gds/LLMOMG/Fasta/bin_60_80.csv", dtype={"seq": str})
test_data_gt80 = pd.read_csv("/projects/intro2gds/LLMOMG/Fasta/bin_gt80.csv", dtype={"seq": str})

# Split by species (90% train, 10% dev)
species = sorted(train_data['Species'].unique())
n_train = 11
train_species = species[:n_train]
dev_species = species[n_train:]

train_split = train_data[train_data['Species'].isin(train_species)].copy()
dev_split = train_data[train_data['Species'].isin(dev_species)].copy()

# Filter orthogroups with >50 sequences in training split only
og_counts = train_split['Orthogroup'].value_counts()
valid_ogs = og_counts[og_counts > 50].index
train_split_filtered = train_split[train_split['Orthogroup'].isin(valid_ogs)].copy()

# Keep only orthogroups shared across all sets
common_ogs = set(train_split_filtered['Orthogroup']) & set(dev_split['Orthogroup']) & set(test_data['Orthogroup'])
print(f"Number of shared orthogroups: {len(common_ogs)}")

train_split = train_split_filtered[train_split_filtered['Orthogroup'].isin(common_ogs)].copy()
dev_split = dev_split[dev_split['Orthogroup'].isin(common_ogs)].copy()
test_data = test_data[test_data['Orthogroup'].isin(common_ogs)].copy()
test_data_lt40 = test_data_lt40[test_data_lt40['Orthogroup'].isin(common_ogs)].copy()
test_data_40_60 = test_data_40_60[test_data_40_60['Orthogroup'].isin(common_ogs)].copy()
test_data_60_80 = test_data_60_80[test_data_60_80['Orthogroup'].isin(common_ogs)].copy()
test_data_gt80 = test_data_gt80[test_data_gt80['Orthogroup'].isin(common_ogs)].copy()

# Label mapping
train_orthogroups = sorted(train_split['Orthogroup'].unique())
orthogroup_to_id = {og: i for i, og in enumerate(train_orthogroups)}
num_classes = len(orthogroup_to_id)

train_split['label'] = train_split['Orthogroup'].map(orthogroup_to_id)
dev_split['label'] = dev_split['Orthogroup'].map(orthogroup_to_id)
test_data['label'] = test_data['Orthogroup'].map(orthogroup_to_id)
test_data_lt40['label'] = test_data_lt40['Orthogroup'].map(orthogroup_to_id)
test_data_40_60['label'] = test_data_40_60['Orthogroup'].map(orthogroup_to_id)
test_data_60_80['label'] = test_data_60_80['Orthogroup'].map(orthogroup_to_id)
test_data_gt80['label'] = test_data_gt80['Orthogroup'].map(orthogroup_to_id)

# Dataset class
class OrthologDataset(Dataset):
    def __init__(self, data, tokenizer, max_length=512):
        self.sequences = data['seq'].tolist()
        self.labels = data['label'].tolist()
        self.tokenizer = tokenizer
        self.max_length = max_length

    def __len__(self):
        return len(self.sequences)

    def __getitem__(self, idx):
        sequence = self.sequences[idx]
        label = self.labels[idx]
        encoding = self.tokenizer(sequence, truncation=True, padding="max_length", max_length=self.max_length, return_tensors="pt")
        return {key: val.squeeze(0) for key, val in encoding.items()}, torch.tensor(label)

# Initialize tokenizer and model
model_name = "facebook/esm2_t6_8M_UR50D"
tokenizer = AutoTokenizer.from_pretrained(model_name)
model = AutoModelForSequenceClassification.from_pretrained(model_name, num_labels=num_classes)

# Data loaders
batch_size = 32
train_loader = DataLoader(OrthologDataset(train_split, tokenizer), batch_size=batch_size, shuffle=True)
dev_loader = DataLoader(OrthologDataset(dev_split, tokenizer), batch_size=batch_size)
test_loader = DataLoader(OrthologDataset(test_data, tokenizer), batch_size=batch_size)
test_loader_lt40 = DataLoader(OrthologDataset(test_data_lt40, tokenizer), batch_size=batch_size)
test_loader_40_60 = DataLoader(OrthologDataset(test_data_40_60, tokenizer), batch_size=batch_size)
test_loader_60_80 = DataLoader(OrthologDataset(test_data_60_80, tokenizer), batch_size=batch_size)
test_loader_gt80 = DataLoader(OrthologDataset(test_data_gt80, tokenizer), batch_size=batch_size)

# Training setup
device = torch.device("cuda" if torch.cuda.is_available() else "cpu")
model.to(device)
optimizer = AdamW(model.parameters(), lr=2e-5)
criterion = CrossEntropyLoss()

# Evaluation function
def evaluate(loader):
    model.eval()
    all_labels, all_preds = [], []
    total_loss = 0
    with torch.no_grad():
        for batch in loader:
            inputs = {k: v.to(device) for k, v in batch[0].items()}
            labels = batch[1].to(device)
            outputs = model(**inputs)
            loss = criterion(outputs.logits, labels)
            total_loss += loss.item()
            preds = torch.argmax(outputs.logits, dim=1)
            all_labels.extend(labels.cpu().numpy())
            all_preds.extend(preds.cpu().numpy())
    avg_loss = total_loss / len(loader)
    accuracy = accuracy_score(all_labels, all_preds)
    precision = precision_score(all_labels, all_preds, average='macro', zero_division=0)
    recall = recall_score(all_labels, all_preds, average='macro', zero_division=0)
    f1 = f1_score(all_labels, all_preds, average='macro', zero_division=0)
    return accuracy, precision, recall, f1, avg_loss, all_labels, all_preds

# Training loop
epochs = 10
train_losses, train_accuracies = [], []
dev_losses, dev_accuracies = [], []
best_dev_f1 = 0.0
best_model_path = "/projects/intro2gds/LLMOMG/Model/ESM2/esm2_15.pt"

for epoch in range(epochs):
    model.train()
    total_loss, correct, total = 0, 0, 0
    for batch in train_loader:
        optimizer.zero_grad()
        inputs = {k: v.to(device) for k, v in batch[0].items()}
        labels = batch[1].to(device)
        outputs = model(**inputs)
        loss = criterion(outputs.logits, labels)
        loss.backward()
        optimizer.step()
        total_loss += loss.item()
        preds = torch.argmax(outputs.logits, dim=1)
        correct += (preds == labels).sum().item()
        total += labels.size(0)

    train_acc = correct / total
    train_losses.append(total_loss / total)
    train_accuracies.append(train_acc)

    dev_acc, _, _, dev_f1, dev_loss, _, _ = evaluate(dev_loader)
    dev_losses.append(dev_loss)
    dev_accuracies.append(dev_acc)

    print(f"Epoch {epoch+1}/{epochs} | Train Loss: {total_loss:.4f}, Acc: {train_acc:.4f} | Dev Loss: {dev_loss:.4f}, Acc: {dev_acc:.4f}")
    if dev_f1 > best_dev_f1:
        best_dev_f1 = dev_f1
        torch.save(model.state_dict(), best_model_path)
        print("Best model saved.")

# Load best model and evaluate
def save_classification_report(labels, preds, file_path):
    report = classification_report(labels, preds, output_dict=True, zero_division=0)
    df = pd.DataFrame(report).transpose()
    os.makedirs(os.path.dirname(file_path), exist_ok=True)
    df.to_csv(file_path)

model.load_state_dict(torch.load(best_model_path))
print("Loaded best model for test evaluation.")

for loader, name in zip([
    test_loader, test_loader_lt40, test_loader_40_60,
    test_loader_60_80, test_loader_gt80
], [
    "test", "test_lt40", "test_40_60", "test_60_80", "test_gt80"
]):
    acc, prec, rec, f1, _, labels, preds = evaluate(loader)
    print(f"{name}: Accuracy={acc:.4f}, Precision={prec:.4f}, Recall={rec:.4f}, F1={f1:.4f}")
    save_classification_report(labels, preds, f"/projects/intro2gds/LLMOMG/Tables/ESM2/classification_report_15_{name}.csv")

# Plot accuracy/loss
plt.figure()
plt.plot(range(1, epochs+1), train_accuracies, label="Train Accuracy")
plt.plot(range(1, epochs+1), dev_accuracies, label="Validation Accuracy")
plt.xlabel("Epoch")
plt.ylabel("Accuracy")
plt.title("Accuracy over Epochs")
plt.legend()
plt.grid(True)
plt.tight_layout()
plt.savefig("/projects/intro2gds/LLMOMG/Figures/ESM2/accuracy_plot_15.png")

plt.figure()
plt.plot(range(1, epochs+1), train_losses, label="Train Loss")
plt.plot(range(1, epochs+1), dev_losses, label="Validation Loss")
plt.xlabel("Epoch")
plt.ylabel("Loss")
plt.title("Loss over Epochs")
plt.legend()
plt.grid(True)
plt.tight_layout()
plt.savefig("/projects/intro2gds/LLMOMG/Figures/ESM2/loss_plot_15.png")
```
