## Supplementary material for "PLM-OMG: Protein Language Model-Based Ortholog Detection for Cross-Species Cell Type Mapping": Scripts: lstm_15.py

```
import os
import random
import pandas as pd
import torch
import torch.nn as nn
from torch.utils.data import Dataset, DataLoader
from sklearn.metrics import accuracy_score, f1_score, recall_score, precision_score, classification_report
from tqdm import tqdm
import matplotlib.pyplot as plt

# Load datasets
train_data = pd.read_csv("/projects/intro2gds/LLMOMG/Fasta/train.csv", dtype={"seq": str})
test_data = pd.read_csv("/projects/intro2gds/LLMOMG/Fasta/test.csv", dtype={"seq": str})
test_data_lt40 = pd.read_csv("/projects/intro2gds/LLMOMG/Fasta/bin_lt40.csv", dtype={"seq": str})
test_data_40_60 = pd.read_csv("/projects/intro2gds/LLMOMG/Fasta/bin_40_60.csv", dtype={"seq": str})
test_data_60_80 = pd.read_csv("/projects/intro2gds/LLMOMG/Fasta/bin_60_80.csv", dtype={"seq": str})
test_data_gt80 = pd.read_csv("/projects/intro2gds/LLMOMG/Fasta/bin_gt80.csv", dtype={"seq": str})

# Split by species (90% train, 10% dev)
species = sorted(train_data['Species'].unique())
n_train = 11
train_species = species[:n_train]
dev_species = species[n_train:]

train_split = train_data[train_data['Species'].isin(train_species)].copy()
dev_split = train_data[train_data['Species'].isin(dev_species)].copy()

# Filter orthogroups with >50 sequences in training split only
og_counts = train_split['Orthogroup'].value_counts()
valid_ogs = og_counts[og_counts > 50].index
train_split_filtered = train_split[train_split['Orthogroup'].isin(valid_ogs)].copy()

# Keep only orthogroups shared across all sets
common_ogs = set(train_split_filtered['Orthogroup']) & set(dev_split['Orthogroup']) & set(test_data['Orthogroup'])
print(f"Number of shared orthogroups: {len(common_ogs)}")

train_split = train_split_filtered[train_split_filtered['Orthogroup'].isin(common_ogs)].copy()
dev_split = dev_split[dev_split['Orthogroup'].isin(common_ogs)].copy()
test_data = test_data[test_data['Orthogroup'].isin(common_ogs)].copy()
test_data_lt40 = test_data_lt40[test_data_lt40['Orthogroup'].isin(common_ogs)].copy()
test_data_40_60 = test_data_40_60[test_data_40_60['Orthogroup'].isin(common_ogs)].copy()
test_data_60_80 = test_data_60_80[test_data_60_80['Orthogroup'].isin(common_ogs)].copy()
test_data_gt80 = test_data_gt80[test_data_gt80['Orthogroup'].isin(common_ogs)].copy()

# Label mapping
train_orthogroups = sorted(train_split['Orthogroup'].unique())
orthogroup_to_id = {og: i for i, og in enumerate(train_orthogroups)}
num_classes = len(orthogroup_to_id)

train_split['label'] = train_split['Orthogroup'].map(orthogroup_to_id)
dev_split['label'] = dev_split['Orthogroup'].map(orthogroup_to_id)
test_data['label'] = test_data['Orthogroup'].map(orthogroup_to_id)
test_data_lt40['label'] = test_data_lt40['Orthogroup'].map(orthogroup_to_id)
test_data_40_60['label'] = test_data_40_60['Orthogroup'].map(orthogroup_to_id)
test_data_60_80['label'] = test_data_60_80['Orthogroup'].map(orthogroup_to_id)
test_data_gt80['label'] = test_data_gt80['Orthogroup'].map(orthogroup_to_id)

# Amino acid vocab
amino_acids = "ACDEFGHIKLMNPQRSTVWY"
aa_to_idx = {aa: idx + 1 for idx, aa in enumerate(amino_acids)}
vocab_size = len(aa_to_idx) + 1

# Dataset
class ProteinDataset(Dataset):
    def __init__(self, data, max_length=512):
        self.sequences = data['seq'].tolist()
        self.labels = data['label'].tolist()
        self.max_length = max_length

    def __len__(self):
        return len(self.sequences)

    def encode_sequence(self, seq):
        encoded = [aa_to_idx.get(aa, 0) for aa in seq[:self.max_length]]
        encoded += [0] * (self.max_length - len(encoded))
        return encoded

    def __getitem__(self, idx):
        return torch.tensor(self.encode_sequence(self.sequences[idx]), dtype=torch.long), \
               torch.tensor(self.labels[idx], dtype=torch.long)

# LSTM model
class ProteinLSTM(nn.Module):
    def __init__(self, vocab_size, embedding_dim, hidden_dim, num_layers, num_classes, dropout=0.5):
        super().__init__()
        self.embedding = nn.Embedding(vocab_size, embedding_dim, padding_idx=0)
        self.lstm = nn.LSTM(embedding_dim, hidden_dim, num_layers=num_layers,
                            batch_first=True, bidirectional=True,
                            dropout=dropout if num_layers > 1 else 0)
        self.dropout = nn.Dropout(dropout)
        self.fc = nn.Linear(hidden_dim * 2, num_classes)

    def forward(self, x):
        x = self.embedding(x)
        lstm_out, _ = self.lstm(x)
        pooled = torch.max(lstm_out, dim=1)[0]
        return self.fc(self.dropout(pooled))

# Metric computation
def calculate_metrics(y_true, y_pred):
    return {
        'accuracy': accuracy_score(y_true, y_pred),
        'precision': precision_score(y_true, y_pred, average='macro', zero_division=0),
        'recall': recall_score(y_true, y_pred, average='macro', zero_division=0),
        'f1': f1_score(y_true, y_pred, average='macro', zero_division=0)
    }

train_losses, dev_losses, train_accuracies, dev_accuracies = [], [], [], []

def train_epoch(model, loader, criterion, optimizer, device):
    model.train()
    total_loss, preds, labels = 0, [], []
    for x, y in tqdm(loader, desc="Training"):
        x, y = x.to(device), y.to(device)
        optimizer.zero_grad()
        out = model(x)
        loss = criterion(out, y)
        loss.backward()
        optimizer.step()
        total_loss += loss.item()
        preds.extend(torch.argmax(out, 1).cpu().numpy())
        labels.extend(y.cpu().numpy())
    metrics = calculate_metrics(labels, preds)
    metrics['loss'] = total_loss / len(loader)
    return metrics

def evaluate(model, loader, criterion, device):
    model.eval()
    total_loss, preds, labels = 0, [], []
    with torch.no_grad():
        for x, y in tqdm(loader, desc="Evaluating"):
            x, y = x.to(device), y.to(device)
            out = model(x)
            loss = criterion(out, y)
            total_loss += loss.item()
            preds.extend(torch.argmax(out, 1).cpu().numpy())
            labels.extend(y.cpu().numpy())
    metrics = calculate_metrics(labels, preds)
    metrics['loss'] = total_loss / len(loader)
    return metrics, labels, preds

def save_classification_report(labels, preds, path):
    report = classification_report(labels, preds, output_dict=True, zero_division=0)
    df = pd.DataFrame(report).transpose()
    os.makedirs(os.path.dirname(path), exist_ok=True)
    df.to_csv(path)

# Training loop
device = torch.device("cuda" if torch.cuda.is_available() else "cpu")
train_loader = DataLoader(ProteinDataset(train_split), batch_size=64, shuffle=True)
dev_loader = DataLoader(ProteinDataset(dev_split), batch_size=64)

model = ProteinLSTM(vocab_size, 128, 256, 2, num_classes, 0.5).to(device)
total_params = sum(p.numel() for p in model.parameters() if p.requires_grad)
print(f"Total trainable parameters: {total_params}")
optimizer = torch.optim.Adam(model.parameters(), lr=0.0001)
criterion = nn.CrossEntropyLoss()

best_f1 = 0
for epoch in range(10):
    print(f"\nEpoch {epoch+1}")
    train_metrics = train_epoch(model, train_loader, criterion, optimizer, device)
    dev_metrics, _, _ = evaluate(model, dev_loader, criterion, device)

    print("Train:", train_metrics)
    print("Dev:  ", dev_metrics)

    train_losses.append(train_metrics['loss'])
    dev_losses.append(dev_metrics['loss'])
    train_accuracies.append(train_metrics['accuracy'])
    dev_accuracies.append(dev_metrics['accuracy'])

    if dev_metrics['f1'] > best_f1:
        best_f1 = dev_metrics['f1']
        torch.save(model.state_dict(), "/projects/intro2gds/LLMOMG/Model/LSTM/LSTM_15.pt")
        print("Saved best model.")

# Evaluate all test sets
test_paths = {
    "original": test_data,
    "lt40": test_data_lt40,
    "40_60": test_data_40_60,
    "60_80": test_data_60_80,
    "gt80": test_data_gt80
}

model.load_state_dict(torch.load("/projects/intro2gds/LLMOMG/Model/LSTM/LSTM_15.pt"))
model.eval()

for name, df in test_paths.items():
    loader = DataLoader(ProteinDataset(df), batch_size=64)
    metrics, labels, preds = evaluate(model, loader, criterion, device)
    print(f"Test ({name}):", metrics)
    save_classification_report(labels, preds, f"/projects/intro2gds/LLMOMG/Tables/LSTM/classification_report_15_{name}.csv")

# Plot accuracy/loss
plt.figure()
plt.plot(range(1, len(train_accuracies)+1), train_accuracies, label="Train Accuracy")
plt.plot(range(1, len(dev_accuracies)+1), dev_accuracies, label="Validation Accuracy")
plt.xlabel("Epoch")
plt.ylabel("Accuracy")
plt.title("Accuracy over Epochs")
plt.legend()
plt.grid(True)
plt.tight_layout()
plt.savefig("/projects/intro2gds/LLMOMG/Figures/LSTM/accuracy_plot_15.png")

plt.figure()
plt.plot(range(1, len(train_losses)+1), train_losses, label="Train Loss")
plt.plot(range(1, len(dev_losses)+1), dev_losses, label="Validation Loss")
plt.xlabel("Epoch")
plt.ylabel("Loss")
plt.title("Loss over Epochs")
plt.legend()
plt.grid(True)
plt.tight_layout()
plt.savefig("/projects/intro2gds/LLMOMG/Figures/LSTM/loss_plot_15.png")
```
