## Supplementary material for "PLM-OMG: Protein Language Model-Based Ortholog Detection for Cross-Species Cell Type Mapping": Scripts: Preprocessing_Analyze.Rmd

````
---
title: "Untitled"
author: "tran chau"
date: "2024-11-22"
output: html_document
---

```{r setup, include=FALSE}
knitr::opts_chunk$set(echo = TRUE)
```

```{r}
species15 = read.csv("/Users/tranchau/Documents/LLM_Orthogroup/OG_15species.csv", row.names = 1)
length(unique(species15$Orthogroup))
length(unique(species15$Species))
Ortho_qc = names(which(table(species15$Orthogroup) > 0))
data = species15[species15$Orthogroup %in% Ortho_qc,]

length(data$seq)


summary(as.numeric(table(species15$Orthogroup)))
length(unique(species15$Orthogroup))
# 67228
length(unique(data$Species))
# 15 species

data = species15
for(i in unique(data$Species)){
  print(i)
  op = length(unique(data[data$Species == i,]$Orthogroup))
  print(op)
}


summary(nchar(data$seq))
```

```{r}
library(ggplot2)

data = species15
orthogroup_counts <- table(data$Orthogroup)

orthogroup_df <- as.data.frame(orthogroup_counts)
colnames(orthogroup_df) <- c("Orthogroup", "Count")


library(scales)

plot_dist <- ggplot(orthogroup_df, aes(x = Count)) +
  geom_histogram(binwidth = 10, color = "black", fill = "blue", alpha = 0.7, boundary = 0, closed = "left") +
  scale_y_continuous(
    trans = "log10",
    breaks = trans_breaks("log10", function(x) 10^x),
    labels = trans_format("log10", math_format(10^.x))
  ) +
  labs(
    x = "Sequences per orthologous group",
    y = expression(log[10]*"(No. of orthologous groups)")
  ) +
  theme_minimal() +
  theme(
    axis.title.x = element_text(size = 20),
    axis.text.x = element_text(size = 20, color = "black"),
    axis.title.y = element_text(size = 20, face = "bold"),
    axis.text.y = element_text(size = 20, color = "black")
  )


pdf("/Volumes/T7 Shield/LLMOMG/Figures/No_Ortholog_distribution_15species.pdf", width = 9, height = 5)
plot_dist
dev.off()
```


```{r}
## For 15 species
library(ggplot2)
library(reshape2)
library(viridis)
data <- as.data.frame(matrix(c(0.36, 0.41, 0.32, 0.30,
                               0.78, 0.82, 0.80, 0.79,
                               0.85, 0.89, 0.85, 0.85,
                               0.86, 0.88, 0.88, 0.86,
                               0.95, 0.95, 0.95, 0.94), ncol=5))
colnames(data) <- c( "ProteinBERT" ,  "LSTM" , "ProGen2" , "ESM2" , "ProtGPT2" )
rownames(data) <- c("accuracy", "precision", "recall", "F1")
data$Metric <- rownames(data)

# Melt the data
data_melted <- melt(data, id.vars = "Metric", variable.name = "Methods", value.name = "Value")


# Rename columns for better readability
colnames(data_melted) <- c("Metric", "Methods", "Value")

# Create the grouped bar chart
plot_metric = ggplot(data_melted, aes(x = Metric, y = Value, fill = Methods)) +
  geom_bar(stat = "identity", position = "dodge", color = "black") +
  geom_text(aes(label = sprintf("%.2f", Value)),  # Format values to 2 decimal places
            position = position_dodge(width = 0.9), vjust = -0.3, size = 4) +
  labs(x = "Metric",
       y = "Value",
       fill = "Methods") +
  theme_minimal() +
  theme(
    axis.text.x = element_text(angle = 0, hjust = 0.5, size =20),
    axis.text.y = element_text(size =20),
    axis.title.x = element_text(size = 25, face = "bold"),
    axis.title.y = element_text(size = 25, face = "bold"),
    text = element_text(size = 15),
    legend.text = element_text(size = 15),
    legend.title = element_text(size = 15, face = "bold"),
    legend.position = "bottom"
  ) +
  scale_fill_manual(values = c("#000004FF", "#56106EFF", "#BB3754FF", "#F98C0AFF", "#FBBC05")) 

pdf("/Volumes/T7 Shield/LLMOMG/Figures/Performance_metric_15species.pdf", width = 9, height = 5)
plot_metric
dev.off()
```

```{r}
# training and validation accuracy
library(ggplot2)

data <- data.frame(
  Epoch = c(1, 2, 3, 4, 5, 6, 7, 8, 9, 10),
  Train = c(0.02, 0.1, 0.2, 0.28, 0.33, 0.37, 0.4, 0.42, 0.43, 0.43), #PROBERT
  Valid = c(0.07, 0.17, 0.24, 0.31, 0.35, 0.38, 0.41, 0.42, 0.43, 0.43) #PROBERT
  #Train = c(0.02, 0.25, 0.55, 0.71, 0.78, 0.83, 0.85, 0.87, 0.89, 0.9), #LSTM
  #Valid = c(0.1, 0.5, 0.69, 0.78, 0.81, 0.84, 0.85, 0.86, 0.88, 0.88) #LSTM
  #Train = c(0.75, 0.94, 0.98, 0.99, 0.99, 0.99, 0.99, 0.99, 1, 1), #PROGEN
  #Valid = c(0.87, 0.89, 0.9, 0.89, 0.89, 0.91, 0.9, 0.9, 0.9, 0.91) #PROGEN
  #Train = c(0.15, 0.34, 0.56, 0.74, 0.86, 0.92, 0.95, 0.97, 0.98, 0.99), #ESM2
  #Valid = c(0.24, 0.43, 0.62, 0.76, 0.84, 0.89, 0.91, 0.92, 0.92, 0.92) #ESM2
  #Train = c(0.57, 0.94, 0.97, 0.99, 0.99, 1, 1, 1, 1, 1), #ProtGPT2
  #Valid = c(0.88, 0.95, 0.95, 0.96, 0.96, 0.96, 0.97, 0.97, 0.97, 0.97) #ProtGPT2
)

library(tidyr)
data_long <- data %>%
  pivot_longer(cols = c(Train, Valid), names_to = "Dataset", values_to = "Accuracy")

# Plot using ggplot2
plot <- ggplot(data_long, aes(x = Epoch, y = Accuracy, color = Dataset)) +
  geom_line(size = 1) +
  geom_point(size = 2) +
  scale_y_continuous(limits = c(0, 1)) +  
  scale_x_continuous(breaks = seq(0, 10, 2)) +
  labs(x = "Epoch", y = "Accuracy", color = "Dataset") +
  theme_minimal(base_size = 14) +
  theme(legend.position = c(0.8, 0.2),
        axis.title.x = element_text(size = 30, face = "bold"),
        axis.text.x = element_text(size = 25, color = "black"),
        axis.title.y = element_text(size = 30, face = "bold"),
        axis.text.y = element_text(size = 25, color = "black"),
        legend.text = element_text(size = 25),
        legend.title = element_text(size = 30, face = "bold")
        )

# Display the plot
print(plot)
pdf("/Volumes/T7 Shield/LLMOMG/Figures/train_valid_PROBERT.pdf", width = 9, height = 5)
plot
dev.off()
```
# Accuracy compare different models in different bins
```{r}
library(fmsb)
library(viridis)

# Prepare your data
data <- as.data.frame(matrix(c(0.19, 0.25, 0.35, 0.54,
                               0.48, 0.67, 0.79, 0.9,
                               0.68, 0.79, 0.86, 0.92,
                               0.63, 0.8, 0.88, 0.93,
                               0.89, 0.93, 0.96, 0.98), ncol = 5))
colnames(data) <- c("ProteinBERT", "LSTM", "ProGen2", "ESM2", "ProtGPT2")
rownames(data) <- c("<40", "40-60", "60-80", ">80")

# Each row is a shape (bin), each column is a model
radar_data <- data

# Add max and min rows
radar_data <- rbind(
  max = rep(1, ncol(radar_data)),
  min = rep(0, ncol(radar_data)),
  radar_data
)
radar_data <- radar_data[, c("ProtGPT2",  "ESM2", "ProGen2", "LSTM", "ProteinBERT" )]
colors <- c("#F7DC6F",  "#58D68D",  "#5DADE2", "#EC7063" )

# Plot radar chart


pdf("/Volumes/T7 Shield/LLMOMG/Figures/mmseq2_acc.pdf", width = 9, height = 9)
radarchart(radar_data,
           axistype = 1,
           seg = 5,  
           pcol = colors,
           pfcol = scales::alpha(colors, 0.08),
           plwd = 2,
           plty = 1,
           cglcol = "gray50",
           cglty = 2,
           cglwd = 0.8,
           axislabcol = "gray30",
           caxislabels = c("0", "0.2", "0.4", "0.6", "0.8", "1"),
           vlcex = 1.5)      
title(main = "Accuracy", cex.main = 1.8, font.main = 2)
legend("bottom", legend = rownames(radar_data)[-c(1, 2)], col = colors, lty = 1, lwd = 2, bty = "n", horiz = TRUE, text.font = 2, cex = 1.3 )
dev.off()
```

# Precision
```{r}
library(fmsb)
library(viridis)

# Prepare your data
data <- as.data.frame(matrix(c(0.41, 0.2, 0.35, 0.39,
                               0.48, 0.67, 0.79, 0.9,
                               0.55, 0.76, 0.86, 0.88,
                               0.5, 0.75, 0.86, 0.86,
                               0.78, 0.9, 0.95, 0.94), ncol = 5))
colnames(data) <- c("ProteinBERT", "LSTM", "ProGen2", "ESM2", "ProtGPT2")
rownames(data) <- c("<40", "40-60", "60-80", ">80")

# Each row is a shape (bin), each column is a model
radar_data <- data

# Add max and min rows
radar_data <- rbind(
  max = rep(1, ncol(radar_data)),
  min = rep(0, ncol(radar_data)),
  radar_data
)
radar_data <- radar_data[, c("ProtGPT2",  "ESM2", "ProGen2", "LSTM", "ProteinBERT" )]
colors <- c("#F7DC6F",  "#58D68D",  "#5DADE2", "#EC7063" )

# Plot radar chart


pdf("/Volumes/T7 Shield/LLMOMG/Figures/mmseq2_precision.pdf", width = 9, height = 9)
radarchart(radar_data,
           axistype = 1,
           seg = 5,  
           pcol = colors,
           pfcol = scales::alpha(colors, 0.08),
           plwd = 2,
           plty = 1,
           cglcol = "gray50",
           cglty = 2,
           cglwd = 0.8,
           axislabcol = "gray30",
           caxislabels = c("0", "0.2", "0.4", "0.6", "0.8", "1"),
           vlcex = 1.5) 
title(main = "Precision", cex.main = 1.8, font.main = 2)
legend("bottom", legend = rownames(radar_data)[-c(1, 2)], col = colors, lty = 1, lwd = 2, bty = "n", horiz = TRUE, text.font = 2, cex = 1.3)
dev.off()
```

# Recall
```{r}
library(fmsb)
library(viridis)

# Prepare your data
data <- as.data.frame(matrix(c(0.12, 0.21, 0.3, 0.38,
                               0.38, 0.63, 0.78, 0.81,
                               0.55, 0.74, 0.85, 0.86,
                               0.51, 0.75, 0.86, 0.85,
                               0.78, 0.89, 0.95, 0.93), ncol = 5))
colnames(data) <- c("ProteinBERT", "LSTM", "ProGen2", "ESM2", "ProtGPT2")
rownames(data) <- c("<40", "40-60", "60-80", ">80")

# Each row is a shape (bin), each column is a model
radar_data <- data

# Add max and min rows
radar_data <- rbind(
  max = rep(1, ncol(radar_data)),
  min = rep(0, ncol(radar_data)),
  radar_data
)
radar_data <- radar_data[, c("ProtGPT2",  "ESM2", "ProGen2", "LSTM", "ProteinBERT" )]
colors <- c("#F7DC6F",  "#58D68D",  "#5DADE2", "#EC7063" )

# Plot radar chart


pdf("/Volumes/T7 Shield/LLMOMG/Figures/mmseq2_recall.pdf", width = 9, height = 9)
radarchart(radar_data,
           axistype = 1,
           seg = 5,  
           pcol = colors,
           pfcol = scales::alpha(colors, 0.08),
           plwd = 2,
           plty = 1,
           cglcol = "gray50",
           cglty = 2,
           cglwd = 0.8,
           axislabcol = "gray30",
           caxislabels = c("0", "0.2", "0.4", "0.6", "0.8", "1"),
           vlcex = 1.5)       
title(main = "Recall", cex.main = 1.8, font.main = 2)
legend("bottom", legend = rownames(radar_data)[-c(1, 2)], col = colors, lty = 1, lwd = 2, bty = "n", horiz = TRUE, text.font = 2, cex = 1.3 )
dev.off()
```

# F1
```{r}
library(fmsb)
library(viridis)

# Prepare your data
data <- as.data.frame(matrix(c(0.1, 0.17, 0.27, 0.33,
                               0.37, 0.6, 0.76, 0.8,
                               0.53, 0.73, 0.84, 0.86,
                               0.49, 0.73, 0.84, 0.85,
                               0.77, 0.88, 0.94, 0.93), ncol = 5))
colnames(data) <- c("ProteinBERT", "LSTM", "ProGen2", "ESM2", "ProtGPT2")
rownames(data) <- c("<40", "40-60", "60-80", ">80")

# Each row is a shape (bin), each column is a model
radar_data <- data

# Add max and min rows
radar_data <- rbind(
  max = rep(1, ncol(radar_data)),
  min = rep(0, ncol(radar_data)),
  radar_data
)
radar_data <- radar_data[, c("ProtGPT2",  "ESM2", "ProGen2", "LSTM", "ProteinBERT" )]
colors <- c("#F7DC6F",  "#58D68D",  "#5DADE2", "#EC7063" )

# Plot radar chart


pdf("/Volumes/T7 Shield/LLMOMG/Figures/mmseq2_f1.pdf", width = 9, height = 9)
radarchart(radar_data,
           axistype = 1,
           seg = 5,  
           pcol = colors,
           pfcol = scales::alpha(colors, 0.08),
           plwd = 2,
           plty = 1,
           cglcol = "gray50",
           cglty = 2,
           cglwd = 0.8,
           axislabcol = "gray30",
           caxislabels = c("0", "0.2", "0.4", "0.6", "0.8", "1"),
           vlcex = 1.5)  
title(main = "F1 Score", cex.main = 1.8, font.main = 2)
legend("bottom", legend = rownames(radar_data)[-c(1, 2)], col = colors, lty = 1, lwd = 2, bty = "n", horiz = TRUE, text.font = 2, cex = 1.3 )
dev.off()
```


```{r}
library(ggplot2)
library(reshape2)
library(dplyr)

# Dummy example: 2 species x 5 models x 4 metrics
accuracy <- c(0.25, 0.53, 0.85, 0.87, 0.88, 0.88, 0.88, 0.91, 0.96, 0.96)
precision <- c(0.18, 0.6, 0.88, 0.91, 0.91, 0.92, 0.87, 0.9, 0.95, 0.96)
recall <- c(0.1, 0.42, 0.84, 0.86, 0.85, 0.87, 0.85, 0.89, 0.96, 0.96)
f1 <- c(0.09, 0.42, 0.85, 0.88, 0.87, 0.89, 0.85, 0.89, 0.95, 0.96)

# Combine into long-format data frame
models <- rep(c("ProteinBERT", "LSTM", "ProGen2", "ESM2", "ProtGPT2"), each = 2)
dataset <- rep(c("Monocots", "Dicots"), times = 5)

df <- data.frame(
  Dataset = dataset,
  Methods = models,
  Accuracy = accuracy,
  Precision = precision,
  Recall = recall,
  F1 = f1
)

# Melt into long format for ggplot
df_long <- melt(df, id.vars = c("Dataset", "Methods"), variable.name = "Metric", value.name = "Value")
df_long$Dataset <- factor(df_long$Dataset, levels = c("Monocots", "Dicots"))

# Plot: Four line plots side by side (one for each metric)
plot_com = ggplot(df_long, aes(x = Dataset, y = Value, color = Methods, group = Methods)) +
  geom_line(linewidth = 1) +
  geom_point(size = 3) +
  facet_wrap(~ Metric, nrow = 1) +
  scale_color_manual(values = c("#F98C0AFF", "#56106EFF", "#000004FF", "#BB3754FF", "#FBBC05")) +
  labs(x = "Dataset", y = "Performance", color = "Methods") +
  theme(
    text = element_text(size = 14),
    axis.title = element_text(size = 16),  
    strip.text = element_text(size = 25, face = "bold"),
    strip.background = element_rect(fill = "gray95", color = "black", linewidth = 0.8),
    axis.text.x = element_text(angle = 0, hjust = 0.5, size = 20),
    axis.text.y = element_text(size = 20),
    axis.title.x = element_text(size = 25, face = "bold"),
    axis.title.y = element_text(size = 25, face = "bold"),
    axis.ticks = element_line(linewidth = 1),
    axis.ticks.length = unit(0.15, "cm"),
    axis.line = element_line(linewidth = 1),
    legend.text = element_text(size = 15),
    legend.title = element_text(size = 15, face = "bold"),
    legend.position = "right"
  )

plot_com
pdf("/Volumes/T7 Shield/LLMOMG/Figures/Dicot_monocot_dataset.pdf", width = 15, height = 5)
plot_com
dev.off()
   
```
````
