## Supplementary material for "PLM-OMG: Protein Language Model-Based Ortholog Detection for Cross-Species Cell Type Mapping": Scripts: progen_15.py

```
# Modified PROGEN-based protein classification code

import os
import pandas as pd
import torch
from torch.utils.data import Dataset, DataLoader
from transformers import AutoTokenizer, AutoModelForCausalLM
from torch.nn import CrossEntropyLoss, Linear
from torch.optim import AdamW
from sklearn.metrics import accuracy_score, precision_score, recall_score, f1_score, classification_report
import matplotlib.pyplot as plt

# Set seed for reproducibility
torch.manual_seed(42)

# Load datasets
train_data = pd.read_csv("/projects/intro2gds/LLMOMG/Fasta/train.csv", dtype={"seq": str})
test_data = pd.read_csv("/projects/intro2gds/LLMOMG/Fasta/test.csv", dtype={"seq": str})
test_data_lt40 = pd.read_csv("/projects/intro2gds/LLMOMG/Fasta/bin_lt40.csv", dtype={"seq": str})
test_data_40_60 = pd.read_csv("/projects/intro2gds/LLMOMG/Fasta/bin_40_60.csv", dtype={"seq": str})
test_data_60_80 = pd.read_csv("/projects/intro2gds/LLMOMG/Fasta/bin_60_80.csv", dtype={"seq": str})
test_data_gt80 = pd.read_csv("/projects/intro2gds/LLMOMG/Fasta/bin_gt80.csv", dtype={"seq": str})

# Split by species (90% train, 10% dev)
species = sorted(train_data['Species'].unique())
n_train = 11
train_species = species[:n_train]
dev_species = species[n_train:]

train_split = train_data[train_data['Species'].isin(train_species)].copy()
dev_split = train_data[train_data['Species'].isin(dev_species)].copy()

# Filter orthogroups with >50 samples in training split only
train_ogs = train_split['Orthogroup'].value_counts()
valid_ogs = train_ogs[train_ogs > 50].index
train_split = train_split[train_split['Orthogroup'].isin(valid_ogs)].copy()

# Keep only shared orthogroups
common_ogs = set(train_split['Orthogroup']) & set(dev_split['Orthogroup']) & set(test_data['Orthogroup'])
train_split = train_split[train_split['Orthogroup'].isin(common_ogs)].copy()
dev_split = dev_split[dev_split['Orthogroup'].isin(common_ogs)].copy()
test_data = test_data[test_data['Orthogroup'].isin(common_ogs)].copy()
test_data_lt40 = test_data_lt40[test_data_lt40['Orthogroup'].isin(common_ogs)].copy()
test_data_40_60 = test_data_40_60[test_data_40_60['Orthogroup'].isin(common_ogs)].copy()
test_data_60_80 = test_data_60_80[test_data_60_80['Orthogroup'].isin(common_ogs)].copy()
test_data_gt80 = test_data_gt80[test_data_gt80['Orthogroup'].isin(common_ogs)].copy()

train_orthogroups = sorted(train_split['Orthogroup'].unique())
orthogroup_to_id = {og: i for i, og in enumerate(train_orthogroups)}

train_split['label'] = train_split['Orthogroup'].map(orthogroup_to_id)
dev_split['label'] = dev_split['Orthogroup'].map(orthogroup_to_id)
test_data['label'] = test_data['Orthogroup'].map(orthogroup_to_id)
test_data_lt40['label'] = test_data_lt40['Orthogroup'].map(orthogroup_to_id)
test_data_40_60['label'] = test_data_40_60['Orthogroup'].map(orthogroup_to_id)
test_data_60_80['label'] = test_data_60_80['Orthogroup'].map(orthogroup_to_id)
test_data_gt80['label'] = test_data_gt80['Orthogroup'].map(orthogroup_to_id)

# Dataset
class OrthologDataset(Dataset):
    def __init__(self, data, tokenizer, max_length=128):
        self.sequences = data['seq'].tolist()
        self.labels = data['label'].tolist()
        self.tokenizer = tokenizer
        self.max_length = max_length

    def __len__(self):
        return len(self.sequences)

    def __getitem__(self, idx):
        sequence = self.sequences[idx]
        label = self.labels[idx]
        inputs = self.tokenizer(sequence, max_length=self.max_length, padding='max_length', truncation=True, return_tensors='pt')
        return {
            'input_ids': inputs['input_ids'].squeeze(0),
            'attention_mask': inputs['attention_mask'].squeeze(0)
        }, torch.tensor(label, dtype=torch.long)

# Model
class ProteinClassifier(torch.nn.Module):
    def __init__(self, model_name, num_classes):
        super().__init__()
        self.base_model = AutoModelForCausalLM.from_pretrained(model_name, trust_remote_code=True)
        emb_dim = self.base_model.transformer.wte.weight.shape[1]
        self.classifier = Linear(emb_dim, num_classes)

    def forward(self, input_ids, attention_mask):
        outputs = self.base_model(input_ids=input_ids, attention_mask=attention_mask, output_hidden_states=True, return_dict=True)
        last_hidden = outputs.hidden_states[-1]
        mask_exp = attention_mask.unsqueeze(-1).expand(last_hidden.size()).float()
        pooled = torch.sum(last_hidden * mask_exp, 1) / torch.clamp(mask_exp.sum(1), min=1e-9)
        return self.classifier(pooled)

# Config
model_name = "hugohrban/progen2-medium"
tokenizer = AutoTokenizer.from_pretrained(model_name, trust_remote_code=True)
tokenizer.pad_token = tokenizer.eos_token
tokenizer.padding_side = 'right'

model = ProteinClassifier(model_name, len(orthogroup_to_id))
print(f"Total trainable parameters: {sum(p.numel() for p in model.parameters() if p.requires_grad)}")

device = torch.device("cuda" if torch.cuda.is_available() else "cpu")
model.to(device)

batch_size = 32
train_loader = DataLoader(OrthologDataset(train_split, tokenizer), batch_size=batch_size, shuffle=True)
dev_loader = DataLoader(OrthologDataset(dev_split, tokenizer), batch_size=batch_size)

optimizer = AdamW(model.parameters(), lr=2e-5)
criterion = CrossEntropyLoss()

# Evaluation function
def evaluate(loader):
    model.eval()
    all_labels, all_preds = [], []
    total_loss = 0
    with torch.no_grad():
        for batch in loader:
            inputs = {k: v.to(device) for k, v in batch[0].items()}
            labels = batch[1].to(device)
            logits = model(**inputs)
            loss = criterion(logits, labels)
            total_loss += loss.item()
            preds = torch.argmax(logits, dim=1)
            all_labels.extend(labels.cpu().numpy())
            all_preds.extend(preds.cpu().numpy())
    acc = accuracy_score(all_labels, all_preds)
    prec = precision_score(all_labels, all_preds, average='macro', zero_division=0)
    rec = recall_score(all_labels, all_preds, average='macro', zero_division=0)
    f1 = f1_score(all_labels, all_preds, average='macro', zero_division=0)
    return acc, prec, rec, f1, total_loss / len(loader), all_labels, all_preds

# Training loop
epochs = 10
best_dev_f1 = 0.0
best_model_path = "/projects/intro2gds/LLMOMG/Model/PROGEN/progen2_15.pt"
train_losses, train_accuracies, dev_losses, dev_accuracies = [], [], [], []

for epoch in range(epochs):
    model.train()
    total_loss, correct, total = 0, 0, 0
    for batch in train_loader:
        optimizer.zero_grad()
        inputs = {k: v.to(device) for k, v in batch[0].items()}
        labels = batch[1].to(device)
        logits = model(**inputs)
        loss = criterion(logits, labels)
        loss.backward()
        optimizer.step()
        total_loss += loss.item()
        preds = torch.argmax(logits, dim=1)
        correct += (preds == labels).sum().item()
        total += labels.size(0)
    train_acc = correct / total
    train_losses.append(total_loss / total)
    train_accuracies.append(train_acc)
    dev_acc, _, _, dev_f1, dev_loss, _, _ = evaluate(dev_loader)
    dev_losses.append(dev_loss)
    dev_accuracies.append(dev_acc)
    print(f"Epoch {epoch+1}/{epochs} | Train Acc: {train_acc:.4f}, Loss: {total_loss:.4f} | Dev Acc: {dev_acc:.4f}, Loss: {dev_loss:.4f}")
    if dev_f1 > best_dev_f1:
        best_dev_f1 = dev_f1
        torch.save(model.state_dict(), best_model_path)
        print("Best model saved.")

# Test set evaluation
def save_classification_report(labels, preds, file_path):
    report = classification_report(labels, preds, output_dict=True, zero_division=0)
    df = pd.DataFrame(report).transpose()
    os.makedirs(os.path.dirname(file_path), exist_ok=True)
    df.to_csv(file_path)

model.load_state_dict(torch.load(best_model_path))
print("Loaded best model for test evaluation.")

test_sets = {
    "original": test_data,
    "lt40": test_data_lt40,
    "40_60": test_data_40_60,
    "60_80": test_data_60_80,
    "gt80": test_data_gt80
}

for name, df in test_sets.items():
    loader = DataLoader(OrthologDataset(df, tokenizer), batch_size=batch_size)
    acc, prec, rec, f1, loss, labels, preds = evaluate(loader)
    print(f"Test ({name}): Accuracy={acc:.4f}, Precision={prec:.4f}, Recall={rec:.4f}, F1={f1:.4f}")
    save_classification_report(labels, preds, f"/projects/intro2gds/LLMOMG/Tables/PROGEN/classification_report_test_15_{name}.csv")

# Plot accuracy/loss
plt.figure()
plt.plot(range(1, epochs+1), train_accuracies, label="Train Accuracy")
plt.plot(range(1, epochs+1), dev_accuracies, label="Validation Accuracy")
plt.xlabel("Epoch")
plt.ylabel("Accuracy")
plt.title("Accuracy over Epochs")
plt.legend()
plt.grid(True)
plt.tight_layout()
plt.savefig("/projects/intro2gds/LLMOMG/Figures/PROGEN/accuracy_plot_15.png")

plt.figure()
plt.plot(range(1, epochs+1), train_losses, label="Train Loss")
plt.plot(range(1, epochs+1), dev_losses, label="Validation Loss")
plt.xlabel("Epoch")
plt.ylabel("Loss")
plt.title("Loss over Epochs")
plt.legend()
plt.grid(True)
plt.tight_layout()
plt.savefig("/projects/intro2gds/LLMOMG/Figures/PROGEN/loss_plot_15.png")
```
