## Supplementary material for "PLM-OMG: Protein Language Model-Based Ortholog Detection for Cross-Species Cell Type Mapping": Scripts: progpt2_15.py

```
import os
import pandas as pd
import torch
from torch.utils.data import Dataset, DataLoader
from transformers import AutoModelForCausalLM, AutoTokenizer
from torch.nn import Linear, CrossEntropyLoss
from torch.optim import AdamW
from sklearn.metrics import accuracy_score, precision_score, recall_score, f1_score, classification_report
from transformers import get_scheduler
from tqdm import tqdm
import matplotlib.pyplot as plt

# Reproducibility
torch.manual_seed(42)

# Load datasets
train_data = pd.read_csv("/projects/intro2gds/LLMOMG/Fasta/train.csv", dtype={"seq": str})
test_data = pd.read_csv("/projects/intro2gds/LLMOMG/Fasta/test.csv", dtype={"seq": str})
test_data_lt40 = pd.read_csv("/projects/intro2gds/LLMOMG/Fasta/bin_lt40.csv", dtype={"seq": str})
test_data_40_60 = pd.read_csv("/projects/intro2gds/LLMOMG/Fasta/bin_40_60.csv", dtype={"seq": str})
test_data_60_80 = pd.read_csv("/projects/intro2gds/LLMOMG/Fasta/bin_60_80.csv", dtype={"seq": str})
test_data_gt80 = pd.read_csv("/projects/intro2gds/LLMOMG/Fasta/bin_gt80.csv", dtype={"seq": str})

# Split by species
species = sorted(train_data['Species'].unique())
n_train = 11
train_species = species[:n_train]
dev_species = species[n_train:]

train_split = train_data[train_data['Species'].isin(train_species)].copy()
dev_split = train_data[train_data['Species'].isin(dev_species)].copy()

# Filter orthogroups with >50 samples in training split
og_counts = train_split['Orthogroup'].value_counts()
valid_ogs = og_counts[og_counts > 50].index
train_split = train_split[train_split['Orthogroup'].isin(valid_ogs)].copy()

# Keep only orthogroups shared with dev and all test sets
common_ogs = set(train_split['Orthogroup']) & set(dev_split['Orthogroup']) & set(test_data['Orthogroup']) 

train_split = train_split[train_split['Orthogroup'].isin(common_ogs)].copy()
dev_split = dev_split[dev_split['Orthogroup'].isin(common_ogs)].copy()
test_data = test_data[test_data['Orthogroup'].isin(common_ogs)].copy()
test_data_lt40 = test_data_lt40[test_data_lt40['Orthogroup'].isin(common_ogs)].copy()
test_data_40_60 = test_data_40_60[test_data_40_60['Orthogroup'].isin(common_ogs)].copy()
test_data_60_80 = test_data_60_80[test_data_60_80['Orthogroup'].isin(common_ogs)].copy()
test_data_gt80 = test_data_gt80[test_data_gt80['Orthogroup'].isin(common_ogs)].copy()

train_orthogroups = sorted(train_split['Orthogroup'].unique())
orthogroup_to_id = {og: i for i, og in enumerate(train_orthogroups)}
num_classes = len(orthogroup_to_id)

# Assign labels
train_split['label'] = train_split['Orthogroup'].map(orthogroup_to_id)
dev_split['label'] = dev_split['Orthogroup'].map(orthogroup_to_id)
test_data['label'] = test_data['Orthogroup'].map(orthogroup_to_id)
test_data_lt40['label'] = test_data_lt40['Orthogroup'].map(orthogroup_to_id)
test_data_40_60['label'] = test_data_40_60['Orthogroup'].map(orthogroup_to_id)
test_data_60_80['label'] = test_data_60_80['Orthogroup'].map(orthogroup_to_id)
test_data_gt80['label'] = test_data_gt80['Orthogroup'].map(orthogroup_to_id)

# Dataset class
class OrthologDataset(Dataset):
    def __init__(self, data, tokenizer, max_length=256):
        self.sequences = data['seq'].tolist()
        self.labels = data['label'].tolist()
        self.tokenizer = tokenizer
        self.max_length = max_length

    def __len__(self):
        return len(self.sequences)

    def __getitem__(self, idx):
        tokens = self.tokenizer(
            self.sequences[idx],
            add_special_tokens=True,
            padding="max_length",
            truncation=True,
            max_length=self.max_length,
            return_tensors="pt"
        )
        return {
            'input_ids': tokens['input_ids'].squeeze(0),
            'attention_mask': tokens['attention_mask'].squeeze(0)
        }, torch.tensor(self.labels[idx])

# Classification model
class ProtGPT2Classifier(torch.nn.Module):
    def __init__(self, base_model, num_labels):
        super().__init__()
        self.base_model = base_model
        self.classifier = Linear(base_model.config.n_embd, num_labels)
        torch.nn.init.kaiming_uniform_(self.classifier.weight, nonlinearity="relu")

    def forward(self, input_ids, attention_mask):
        outputs = self.base_model(input_ids=input_ids, attention_mask=attention_mask, output_hidden_states=True)
        hidden_states = outputs.hidden_states[-1]
        pooled_output = hidden_states.mean(dim=1)
        return self.classifier(pooled_output)

# Instantiate model and training utilities
tokenizer = AutoTokenizer.from_pretrained("nferruz/ProtGPT2")
if tokenizer.pad_token is None:
    tokenizer.pad_token = tokenizer.eos_token

base_model = AutoModelForCausalLM.from_pretrained("nferruz/ProtGPT2")
model = ProtGPT2Classifier(base_model, num_classes)
device = torch.device("cuda" if torch.cuda.is_available() else "cpu")
model.to(device)

# Dataloaders
batch_size = 16
train_loader = DataLoader(OrthologDataset(train_split, tokenizer), batch_size=batch_size, shuffle=True)
dev_loader = DataLoader(OrthologDataset(dev_split, tokenizer), batch_size=batch_size)

# Optimizer and Scheduler
optimizer = AdamW(model.parameters(), lr=5e-5)
num_training_steps = len(train_loader) * 10
scheduler = get_scheduler("linear", optimizer=optimizer, num_warmup_steps=int(0.1 * num_training_steps), num_training_steps=num_training_steps)

# Loss function with class weights
label_counts = train_split['label'].value_counts()
class_weights = torch.tensor([1.0 / label_counts.get(i, 1) for i in range(num_classes)], dtype=torch.float).to(device)
criterion = CrossEntropyLoss(weight=class_weights)

# Evaluation function
def evaluate(loader):
    model.eval()
    all_labels, all_preds = [], []
    total_loss = 0
    with torch.no_grad():
        for batch in loader:
            inputs = {k: v.to(device) for k, v in batch[0].items()}
            labels = batch[1].to(device)
            logits = model(**inputs)
            loss = criterion(logits, labels)
            total_loss += loss.item()
            preds = torch.argmax(logits, dim=1)
            all_labels.extend(labels.cpu().numpy())
            all_preds.extend(preds.cpu().numpy())
    acc = accuracy_score(all_labels, all_preds)
    prec = precision_score(all_labels, all_preds, average='macro', zero_division=0)
    rec = recall_score(all_labels, all_preds, average='macro', zero_division=0)
    f1 = f1_score(all_labels, all_preds, average='macro', zero_division=0)
    return acc, prec, rec, f1, total_loss / len(loader), all_labels, all_preds

# Training loop
train_losses, train_accuracies = [], []
dev_losses, dev_accuracies = [], []
best_dev_f1 = 0.0
best_model_path = "/projects/intro2gds/LLMOMG/Model/PROGPT2/protgpt2_15N.pt"

for epoch in range(10):
    model.train()
    total_loss, correct, total = 0, 0, 0
    for batch in train_loader:
        optimizer.zero_grad()
        inputs = {k: v.to(device) for k, v in batch[0].items()}
        labels = batch[1].to(device)
        logits = model(**inputs)
        loss = criterion(logits, labels)
        loss.backward()
        optimizer.step()
        scheduler.step()
        total_loss += loss.item()
        preds = torch.argmax(logits, dim=1)
        correct += (preds == labels).sum().item()
        total += labels.size(0)
    train_acc = correct / total
    train_losses.append(total_loss / total)
    train_accuracies.append(train_acc)

    dev_acc, _, _, dev_f1, dev_loss, _, _ = evaluate(dev_loader)
    dev_losses.append(dev_loss)
    dev_accuracies.append(dev_acc)

    print(f"Epoch {epoch+1} | Train Acc: {train_acc:.4f} | Dev Acc: {dev_acc:.4f} | Dev F1: {dev_f1:.4f}")
    if dev_f1 > best_dev_f1:
        best_dev_f1 = dev_f1
        torch.save(model.state_dict(), best_model_path)
        print("Best model saved.")

# Test evaluation
model.load_state_dict(torch.load(best_model_path))
print("Loaded best model for test evaluation.")

test_sets = {
    "test": test_data,
    "lt40": test_data_lt40,
    "40_60": test_data_40_60,
    "60_80": test_data_60_80,
    "gt80": test_data_gt80
}

def save_classification_report(labels, preds, file_path):
    report = classification_report(labels, preds, output_dict=True, zero_division=0)
    df = pd.DataFrame(report).transpose()
    os.makedirs(os.path.dirname(file_path), exist_ok=True)
    df.to_csv(file_path)

for name, df in test_sets.items():
    loader = DataLoader(OrthologDataset(df, tokenizer), batch_size=batch_size)
    acc, prec, rec, f1, _, labels, preds = evaluate(loader)
    print(f"{name.upper()} Test: Accuracy={acc:.4f}, Precision={prec:.4f}, Recall={rec:.4f}, F1={f1:.4f}")
    save_path = f"/projects/intro2gds/LLMOMG/Tables/PROGPT2/classification_report_test_15N_{name}.csv"
    save_classification_report(labels, preds, save_path)

# Plot Accuracy and Loss
plt.figure()
plt.plot(range(1, 6), train_accuracies, label="Train Accuracy")
plt.plot(range(1, 6), dev_accuracies, label="Validation Accuracy")
plt.xlabel("Epoch")
plt.ylabel("Accuracy")
plt.title("Accuracy over Epochs")
plt.legend()
plt.grid(True)
plt.tight_layout()
plt.savefig("/projects/intro2gds/LLMOMG/Figures/PROGPT2/accuracy_plot_15N.png")

plt.figure()
plt.plot(range(1, 6), train_losses, label="Train Loss")
plt.plot(range(1, 6), dev_losses, label="Validation Loss")
plt.xlabel("Epoch")
plt.ylabel("Loss")
plt.title("Loss over Epochs")
plt.legend()
plt.grid(True)
plt.tight_layout()
plt.savefig("/projects/intro2gds/LLMOMG/Figures/PROGPT2/loss_plot_15N.png")
```
