## Supplementary material for "PLM-OMG: Protein Language Model-Based Ortholog Detection for Cross-Species Cell Type Mapping": Scripts: proteinbert_15_100.py

```
import os
import pandas as pd
import random
import torch
from torch.utils.data import Dataset, DataLoader
from transformers import AutoTokenizer, BertForSequenceClassification
from torch.nn import CrossEntropyLoss
from torch.optim import AdamW
from transformers import get_scheduler
from sklearn.metrics import accuracy_score, precision_score, recall_score, f1_score, classification_report
from tqdm import tqdm
import numpy as np
import matplotlib.pyplot as plt

# Set seed for reproducibility
torch.manual_seed(42)

# Dataset Class
class ProteinDataset(Dataset):
    def __init__(self, data, tokenizer, max_length=512):
        self.sequences = data['seq'].tolist()
        self.labels = data['label'].tolist()
        self.tokenizer = tokenizer
        self.max_length = max_length

    def __len__(self):
        return len(self.sequences)

    def __getitem__(self, idx):
        sequence = self.sequences[idx]
        label = self.labels[idx]
        encoding = self.tokenizer(
            sequence,
            truncation=True,
            padding="max_length",
            max_length=self.max_length,
            return_tensors="pt"
        )
        return {key: val.squeeze(0) for key, val in encoding.items()}, torch.tensor(label)

# Training function
def train_epoch(model, dataloader, optimizer, scheduler, criterion, device):
    model.train()
    total_loss = 0
    all_preds = []
    all_labels = []

    progress_bar = tqdm(dataloader, desc="Training")
    for batch in progress_bar:
        inputs = {key: val.to(device) for key, val in batch[0].items()}
        labels = batch[1].to(device)

        outputs = model(**inputs)
        loss = criterion(outputs.logits, labels)
        loss.backward()

        torch.nn.utils.clip_grad_norm_(model.parameters(), max_norm=1.0)
        optimizer.step()
        scheduler.step()
        optimizer.zero_grad()

        total_loss += loss.item()
        preds = torch.argmax(outputs.logits, dim=1)
        all_preds.extend(preds.cpu().numpy())
        all_labels.extend(labels.cpu().numpy())

        progress_bar.set_postfix({'loss': f"{loss.item():.4f}"})

    metrics = {
        'loss': total_loss / len(dataloader),
        'accuracy': accuracy_score(all_labels, all_preds),
        'macro_f1': f1_score(all_labels, all_preds, average='macro', zero_division=0),
        'weighted_f1': f1_score(all_labels, all_preds, average='weighted', zero_division=0)
    }
    return metrics

# Evaluation function
def evaluate(model, dataloader, criterion, device):
    model.eval()
    total_loss = 0
    all_preds = []
    all_labels = []

    with torch.no_grad():
        for batch in tqdm(dataloader, desc="Evaluating"):
            inputs = {key: val.to(device) for key, val in batch[0].items()}
            labels = batch[1].to(device)

            outputs = model(**inputs)
            loss = criterion(outputs.logits, labels)

            total_loss += loss.item()
            preds = torch.argmax(outputs.logits, dim=1)
            all_preds.extend(preds.cpu().numpy())
            all_labels.extend(labels.cpu().numpy())

    metrics = {
        'loss': total_loss / len(dataloader),
        'accuracy': accuracy_score(all_labels, all_preds),
        'macro_f1': f1_score(all_labels, all_preds, average='macro', zero_division=0),
        'weighted_f1': f1_score(all_labels, all_preds, average='weighted', zero_division=0)
    }
    return metrics, all_labels, all_preds

# Load datasets
train_data = pd.read_csv("/projects/intro2gds/LLMOMG/Fasta/train.csv", dtype={"seq": str})
test_data = pd.read_csv("/projects/intro2gds/LLMOMG/Fasta/test.csv", dtype={"seq": str})
test_data_lt40 = pd.read_csv("/projects/intro2gds/LLMOMG/Fasta/bin_lt40.csv", dtype={"seq": str})
test_data_40_60 = pd.read_csv("/projects/intro2gds/LLMOMG/Fasta/bin_40_60.csv", dtype={"seq": str})
test_data_60_80 = pd.read_csv("/projects/intro2gds/LLMOMG/Fasta/bin_60_80.csv", dtype={"seq": str})
test_data_gt80 = pd.read_csv("/projects/intro2gds/LLMOMG/Fasta/bin_gt80.csv", dtype={"seq": str})

# Split by species
species = sorted(train_data['Species'].unique())
n_train = 11
train_species = species[:n_train]
dev_species = species[n_train:]

train_split = train_data[train_data['Species'].isin(train_species)].copy()
dev_split = train_data[train_data['Species'].isin(dev_species)].copy()

# Filter orthogroups with >100 samples in training split only
train_ogs = train_split['Orthogroup'].value_counts()
valid_ogs = train_ogs[train_ogs > 100].index
train_split = train_split[train_split['Orthogroup'].isin(valid_ogs)].copy()

# Keep only shared orthogroups
common_ogs = set(train_split['Orthogroup']) & set(dev_split['Orthogroup']) & set(test_data['Orthogroup'])
train_split = train_split[train_split['Orthogroup'].isin(common_ogs)].copy()
dev_split = dev_split[dev_split['Orthogroup'].isin(common_ogs)].copy()
test_data = test_data[test_data['Orthogroup'].isin(common_ogs)].copy()
test_data_lt40 = test_data_lt40[test_data_lt40['Orthogroup'].isin(common_ogs)].copy()
test_data_40_60 = test_data_40_60[test_data_40_60['Orthogroup'].isin(common_ogs)].copy()
test_data_60_80 = test_data_60_80[test_data_60_80['Orthogroup'].isin(common_ogs)].copy()
test_data_gt80 = test_data_gt80[test_data_gt80['Orthogroup'].isin(common_ogs)].copy()

train_orthogroups = sorted(train_split['Orthogroup'].unique())
orthogroup_to_id = {og: i for i, og in enumerate(train_orthogroups)}

train_split['label'] = train_split['Orthogroup'].map(orthogroup_to_id)
dev_split['label'] = dev_split['Orthogroup'].map(orthogroup_to_id)
test_data['label'] = test_data['Orthogroup'].map(orthogroup_to_id)
test_data_lt40['label'] = test_data_lt40['Orthogroup'].map(orthogroup_to_id)
test_data_40_60['label'] = test_data_40_60['Orthogroup'].map(orthogroup_to_id)
test_data_60_80['label'] = test_data_60_80['Orthogroup'].map(orthogroup_to_id)
test_data_gt80['label'] = test_data_gt80['Orthogroup'].map(orthogroup_to_id)

# Tokenizer and Model
model_name = "unikei/bert-base-proteins"
tokenizer = AutoTokenizer.from_pretrained(model_name)
model = BertForSequenceClassification.from_pretrained(model_name, num_labels=len(orthogroup_to_id), ignore_mismatched_sizes=True)
model.classifier = torch.nn.Linear(model.config.hidden_size, len(orthogroup_to_id))
device = torch.device("cuda" if torch.cuda.is_available() else "cpu")
model.to(device)

# Parameter count
print(f"Total parameters: {sum(p.numel() for p in model.parameters()):,}")
print(f"Trainable parameters: {sum(p.numel() for p in model.parameters() if p.requires_grad):,}")

# Dataloaders
batch_size = 32
train_loader = DataLoader(ProteinDataset(train_split, tokenizer), batch_size=batch_size, shuffle=True)
dev_loader = DataLoader(ProteinDataset(dev_split, tokenizer), batch_size=batch_size)

# Optimizer, Scheduler, Loss
optimizer = AdamW(model.parameters(), lr=1e-6)
epochs = 10
num_training_steps = len(train_loader) * epochs
scheduler = get_scheduler("linear", optimizer=optimizer, num_warmup_steps=int(0.1 * num_training_steps), num_training_steps=num_training_steps)
criterion = CrossEntropyLoss()

# Training Loop
train_accuracies, dev_accuracies, train_losses, dev_losses = [], [], [], []
best_dev_f1 = 0.0
best_model_path = "/projects/intro2gds/LLMOMG/Model/PROBERT/proteinbert_15_100.pt"

for epoch in range(epochs):
    print(f"\nEpoch {epoch + 1}/{epochs}")
    train_metrics = train_epoch(model, train_loader, optimizer, scheduler, criterion, device)
    dev_metrics, _, _ = evaluate(model, dev_loader, criterion, device)

    print("Training metrics:", train_metrics)
    print("Validation metrics:", dev_metrics)

    train_accuracies.append(train_metrics['accuracy'])
    dev_accuracies.append(dev_metrics['accuracy'])
    train_losses.append(train_metrics['loss'])
    dev_losses.append(dev_metrics['loss'])

    if dev_metrics['macro_f1'] > best_dev_f1:
        best_dev_f1 = dev_metrics['macro_f1']
        torch.save(model.state_dict(), best_model_path)
        print(f"New best model saved with F1: {best_dev_f1:.4f}")

# Final evaluation
model.load_state_dict(torch.load(best_model_path))
print("Loaded best model for test evaluation.")

def save_classification_report(labels, preds, file_path):
    report = classification_report(labels, preds, output_dict=True, zero_division=0)
    df = pd.DataFrame(report).transpose()
    os.makedirs(os.path.dirname(file_path), exist_ok=True)
    df.to_csv(file_path)

# Test evaluations
test_sets = {
    "original": test_data,
    "lt40": test_data_lt40,
    "40_60": test_data_40_60,
    "60_80": test_data_60_80,
    "gt80": test_data_gt80
}

for name, df in test_sets.items():
    test_loader = DataLoader(ProteinDataset(df, tokenizer), batch_size=batch_size)
    _, labels, preds = evaluate(model, test_loader, criterion, device)
    acc = accuracy_score(labels, preds)
    prec = precision_score(labels, preds, average='macro', zero_division=0)
    rec = recall_score(labels, preds, average='macro', zero_division=0)
    f1 = f1_score(labels, preds, average='macro', zero_division=0)
    print(f"{name.upper()} Test: Accuracy={acc:.4f}, Precision={prec:.4f}, Recall={rec:.4f}, F1={f1:.4f}")
    save_classification_report(labels, preds, f"/projects/intro2gds/LLMOMG/Tables/PROBERT/classification_report_test_15_100_{name}.csv")

# Plot Accuracy and Loss Curves
plt.figure()
plt.plot(range(1, epochs+1), train_accuracies, label="Train Accuracy")
plt.plot(range(1, epochs+1), dev_accuracies, label="Validation Accuracy")
plt.xlabel("Epoch")
plt.ylabel("Accuracy")
plt.title("Accuracy over Epochs")
plt.legend()
plt.grid(True)
plt.tight_layout()
plt.savefig("/projects/intro2gds/LLMOMG/Figures/PROBERT/accuracy_plot_15_100.png")

plt.figure()
plt.plot(range(1, epochs+1), train_losses, label="Train Loss")
plt.plot(range(1, epochs+1), dev_losses, label="Validation Loss")
plt.xlabel("Epoch")
plt.ylabel("Loss")
plt.title("Loss over Epochs")
plt.legend()
plt.grid(True)
plt.tight_layout()
plt.savefig("/projects/intro2gds/LLMOMG/Figures/PROBERT/loss_plot_15_100.png")
```
